# *Aotus nancymaae* model predicts human immune response to the placental malaria vaccine candidate VAR2CSA

**DOI:** 10.1101/2022.06.24.497389

**Authors:** Justin Doritchamou, Morten A. Nielsen, Arnaud Chêne, Nicola K. Viebig, Lynn E. Lambert, Adam F. Sander, Jean-Philippe Semblat, Sophia Hundt, Sachy Orr-Gonzalez, Christoph M. Janitzek, Alicia J. Spiegel, Stine B. Clemmensen, Marvin L. Thomas, Martha C. Nason, Maryonne Snow-Smith, Emma K. Barnafo, Joseph Shiloach, Beth B. Chen, Steven Nadakal, Kendrick Highsmith, Tarik Ouahes, Solomon Conteh, Ankur Sharma, Holly Torano, Brandi Butler, Karine Reiter, Kelly M. Rausch, Puthupparampil V. Scaria, Charles Anderson, David L. Narum, Ali Salanti, Michal Fried, Thor G. Theander, Benoit Gamain, Patrick E. Duffy

## Abstract

Placental malaria vaccines (PMV) are being developed to prevent severe sequelae of placental malaria (PM) in pregnant women and their offspring. The leading candidate vaccine antigen VAR2CSA mediates parasite binding to placental receptor chondroitin sulfate A (CSA). Despite promising results in small animal studies, recent human trials of the first two PMV candidates (PAMVAC and PRIMVAC) generated limited cross-reactivity and cross-inhibitory activity to heterologous parasites. Here, we immunized *Aotus nancymaae* monkeys with three PMV candidates (PAMVAC, PRIMVAC and ID1-ID2a_M1010) adjuvanted with Alhydrogel®, and exploited the model to investigate boosting of functional vaccine responses during PM episodes as well as with nanoparticle antigens. PMV candidates induced high levels of antigen-specific IgG with significant cross-reactivity across PMV antigens by ELISA. Conversely, PMV antibodies recognized native VAR2CSA and blocked CSA-adhesion of only homologous parasites and not heterologous parasites. PM episodes did not significantly boost VAR2CSA antibody levels or serum functional activity; nanoparticle and monomer antigens alike boosted serum reactivity but not functional activities. Overall, PMV candidates induced functional antibodies with limited heterologous activity in *Aotus* monkeys, similar to responses reported in humans. The *Aotus* model appears suitable for preclinical down-selection of PMV candidates and assessment of antibody boosting by PM episodes.

**Research in Context:** *Evidence before this study:* The *Plasmodium falciparum* erythrocyte membrane protein VAR2CSA is the leading vaccine candidate antigen to protect pregnant women against placental malaria (PM), which causes serious adverse pregnancy outcomes particularly in first-time mothers living in malaria-endemic areas. Two VAR2CSA-based vaccines (PAMVAC and PRIMVAC) induced strong heterologous functional antibodies in small animals, but induced antibodies with limited cross-inhibitory functional activity in human clinical trials. These observations highlighted the need to establish new animal models that could better recapitulate human pathogenesis and immunity. In ongoing development of a nonhuman primate model for PM, we established an *Aotus nancymaae* model susceptible to *P. falciparum* infection during pregnancy that reproduces all the immunoparasitological and histological features of human PM. In this study, we explore the new *Aotus* model as a platform for evaluating PM vaccine (PMV) immunogenicity and for boosting of vaccine responses during PM episodes.

*Added value of this study:* In this manuscript, we demonstrate that PMV (including PAMVAC and PRIMVAC) are immunogenic in *Aotus* monkeys, inducing antibodies with mainly homologous and little heterologous functional activity, as seen in humans but contrary to preclinical reports on these vaccines in small animals.

*Implications of all the available evidence:* Our findings suggest *Aotus* is a suitable model to assess immunogenicity of VAR2CSA-derived vaccines, in contrast to small animal models. PMV data from human trials and *Aotus* monkeys suggest that improvements to current VAR2CSA immunogens and/or adjuvants are needed to enhance protective antibody responses, as are studies that evaluate the potential for natural infection to boost vaccine antibody in pregnancy. Therefore, the *Aotus* PM model may be useful to assess second-generation PMVs seeking to increase strain-transcending activity and to prioritize these for further clinical development.

## Introduction

Pregnant women living in malaria-endemic areas become more susceptible to *Plasmodium* infection despite pre-existing immunity to malaria acquired during childhood (1). *P. falciparum* malaria during pregnancy leads to placental malaria (PM), which is characterized by the sequestration of chondroitin sulfate A (CSA)-binding *P. falciparum-*infected erythrocytes (IE) in the placenta and is associated with adverse pregnancy outcomes (2-4). Susceptibility to PM is more pronounced in first-time mothers and decreases over successive pregnancies, as women acquire functional anti-adhesion antibodies against CSA-binding IE (5, 6). These observations suggest that resistance to PM can be conferred to pregnant women through vaccination by inducing protective antibodies targeting the surface of placental IE (6).

The placenta-sequestering *P. falciparum* IE surface displays VAR2CSA, a member of the *P. falciparum* erythrocyte membrane protein 1 (*Pf*EMP1) family and the main parasite ligand mediating IE adhesion to the key placental receptor CSA (7-10). The large size of VAR2CSA stymies efforts to manufacture full-length recombinant protein, and studies of VAR2CSA domains or fragments that bind CSA have informed the design of constructs that induce anti-adhesion antibodies in small animals (mouse, rat and rabbit) (11-19). Among these VAR2CSA constructs, PAMVAC and PRIMVAC, based on partially overlapping N-terminal fragments of VAR2CSA, have recently been tested in phase I clinical trials in Europe and Africa (ClinicalTrials.gov, NCT02647489 and NCT02658253). In rodents, PAMVAC and PRIMVAC induced cross-reactive binding-inhibitory antisera with activity similar to that detected in PM-resistant multigravid women (12, 17, 20).

First-in-human reports from clinical trials demonstrated that both PAMVAC and PRIMVAC vaccines were safe, well-tolerated, immunogenic, and induced functional antibodies in malaria-naïve and malaria-exposed women (21, 22). However, the level of anti-adhesion activity of antibodies induced in volunteers appeared lower compared to that found when PAMVAC and PRIMVAC were tested in rodents with the same adjuvants (12, 17, 20-22), with limited cross-strain activity, suggesting that these VAR2CSA-based vaccines could be improved or combined to generate broader protection against PM. These observations also highlight the importance of relevant animal models that could better predict human immune responses to VAR2CSA antigens.

Here, we evaluate a nonhuman primate (NHP) model for assessing VAR2CSA-based PM vaccines (PMV). Many studies have indicated the *Aotus nancymaae* monkey model to be useful for supporting malaria vaccine development, since it is susceptible to *P. falciparum* infection, and shows antibody profiles similar to humans following a malaria infection (23-25). Furthermore, we have recently established an *Aotus* model for PM that recapitulates key features of malaria infection and immunity in pregnant women, including placental sequestration, selective binding to CSA by placental parasites, and the acquisition of heterologous functional antibodies over successive pregnancies (Sharma et al., JID 2022). Overall, this study provides evidence that PMV immunogenicity in the new *Aotus* model is similar to that of human responses.

## Methods

### Study design

A blinded, multi-phase study was designed to test three PMV candidates (**Fig. 1A**) using a newly established *Aotus* monkey model for PM (Sharma et al., JID 2022). In the first phase of this study, 40 naïve female *Aotus nancymaae* were randomized to receive the VAR2CSA-based vaccine candidates PAMVAC (n = 9), PRIMVAC (n = 9) and ID1-ID2a_M1010 (n = 9) as well as a transmission-blocking vaccine candidate Pfs25 (n = 13) to serve as a control. Animals were immunized in 2 cohorts of 20 monkeys/cohort with ∼5 months interval between both cohorts. For the initial 20 animals (Cohort 1), we assigned 4 monkeys to each PMV vaccine group and 8 monkeys to the control group. For the second 20 animals (Cohort 2), 5 animals per group were assigned to improve the overall power. Vaccines were formulated on Alhydrogel® and each monkey received 3 immunizations at 4-week intervals (day (D) 0, D28, D56), comprised of 50 µg of vaccine by intramuscular injection in the thigh, alternating legs. Blood samples were collected at D0 and D70 to assess induction of functional antibodies following vaccination.

**Figure 1.**
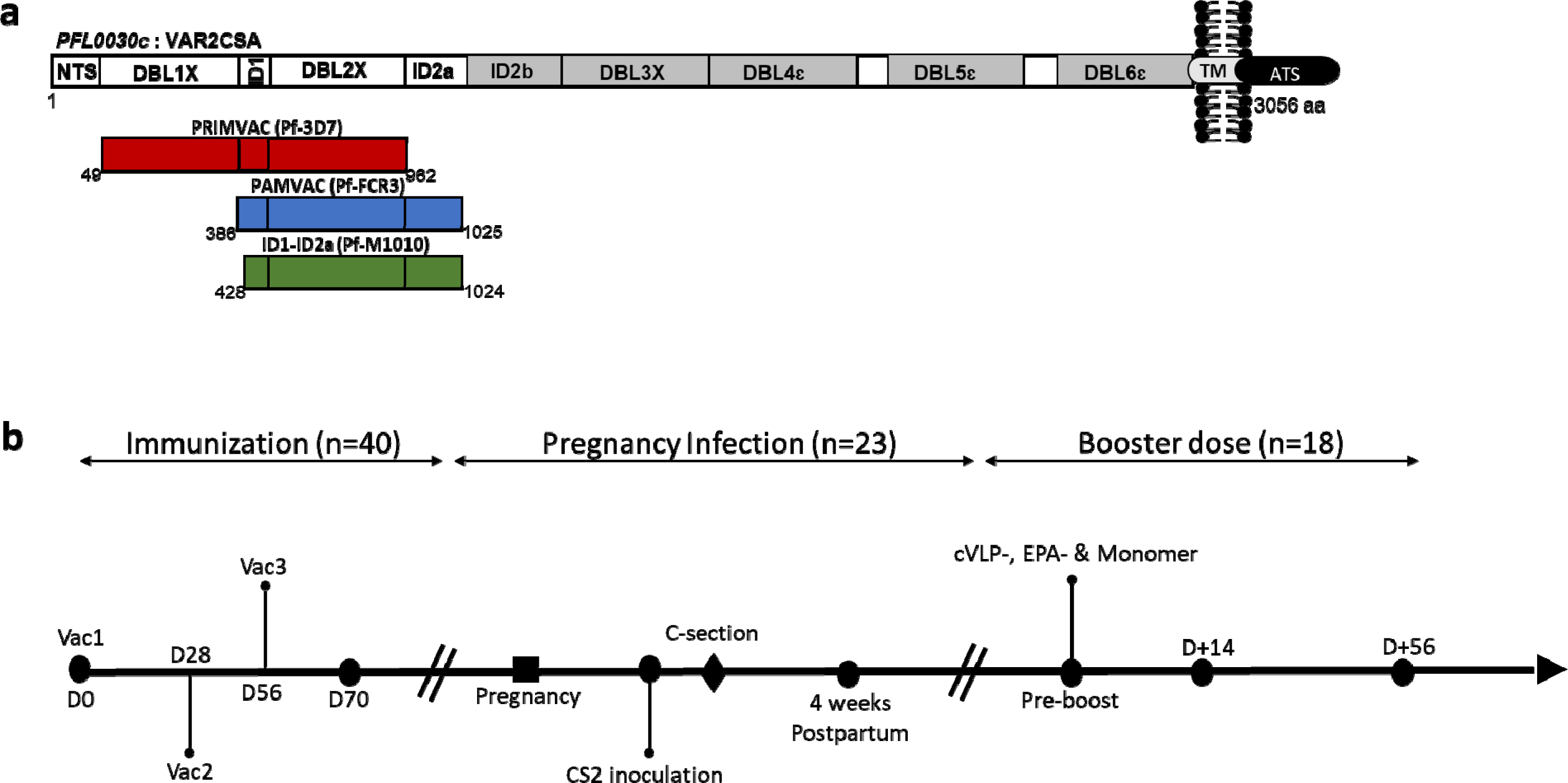
PMV constructs and study design. a) Illustration of three PMV constructs of VAR2CSA used in this study. Boundaries of each construct are shown as amino acid residues from the corresponding VAR2CSA allele (FCR3 and 3D7) and M1010 composite sequence. b) Flow chart of the study highlighting different stages of the study and number of animals involved in each stage. Timeline indicates three vaccinations (Vac1, Vac2 and Vac3) with PMV and Pfs25 antigens. C-section = Cesarean section, cVLP = capsid-based virus like particle; EPA = mutagenized *Pseudomonas aeruginosa* exoprotein A.

In the second phase of the study, 23 immunized monkeys (6 with PAMVAC, 5 with PRIMVAC, 4 with ID1-ID2a_M1010 and 8 with Pfs25) that became pregnant were infected with *P. falciparum* CS2 clone (number of IEs ranging from ∼1·2 to 2·5 x 10^7^). For this infection in pregnant monkeys, stocks of CS2 inoculum for this study were obtained after sub-passing CS2 parasite (in human blood) into a male monkey to collect infected blood for CS2 culture in *Aotus* blood. Parasites were then maintained in culture over 4 to 6 weeks and stocks of CS2 inoculum were made at 1% parasite in *Aotus* blood and frozen. Two lots of CS2 inoculums with similar parasite binding phenotype (predominantly binding to CD36 receptor) were used in this work (**Table S1**). CS2 infections in pregnant monkeys were performed with thawed parasites inoculated in the saphenous vein. Infected animals were observed twice a day and monitored daily with parasitemia data collected from day 3 until animals were cured. *P. falciparum* infection was confirmed by PCR detection on peripheral blood 3 days after CS2 inoculation (see **Data File S2** for detailed parasitological data and other clinically relevant information). The infected pregnant monkeys were asymptomatic during this brief CS2 infection, and all animals were cured with mefloquine after end of pregnancy. As full normal gestation is ∼19 weeks (27), infection was initiated at ∼17 weeks to allow for 7 days infection, and C-section performed at ∼18 weeks to ensure placental sample collection. Samples were collected on the day of CS2 inoculation and 4 weeks postpartum to investigate whether vaccine-induced antibodies can be boosted or enhanced by a malaria episode during pregnancy. Although the study was not designed to compare parasite densities between groups, thin blood smear was prepared from peripheral and placental blood at delivery where available. Of note 14 monkeys completed pregnancy with scheduled C-section whereas 9 animals delivered naturally and 2 had stillbirth prior to C-section precluding collection of placenta samples.

The third phase of the study assessed PRIMVAC and PAMVAC vaccine-induced antibodies following a booster dose of cVLP modified to present the PAMVAC antigen or monomer PAMVAC. The vaccine antigen was coupled to preformed cVLP through a split protein covalent interaction to form PAMVAC_cVLP_ nanoparticle as previously described (28, 29). Similar assessment was conducted in monkeys immunized with the ID1-ID2a_M1010 candidate using the ExoProtein A (EPA) chemically conjugated with ID1-ID2a_M1010 to generate ID1-ID2a_M1010_EPA_ nanoparticle as previously described (30). Both PAMVAC_cVLP_ and ID1-ID2a_M1010_EPA_ form nanoparticles. For this phase, 13 monkeys immunized with either PAMVAC (n = 7) or PRIMVAC (n = 6) received either the nanoparticle or monomer forms of PAMVAC, while 5 monkeys immunized with ID1-ID2a_M1010 (n = 5) received either the nanoparticle or monomer forms of ID1-ID2a_M1010. Thus, only PRIMVAC-immunized monkeys received a heterologous vaccine (PAMVAC). Pre-bleed blood samples as well as those collected 14- and 56-days post-booster dose injection were used to investigate whether a boost with PMV nanoparticles can enhance functional activity of PMV vaccine-induced antibodies.

The collected monkey sera from the different study timepoints were assayed for antibody reactivity to PMV and Pfs25 antigens by ELISA. Plasma antibodies were tested for IEs surface reactivity by Flow cytometry and CSA-binding inhibition activity using CS2 (that carries the same VAR2CSA sequence as FCR3 variant used in PAMVAC) and NF54 (that displays the same VAR2CSA sequence as 3D7 variant used in PRIMVAC) isolates. A clonal M1010 isolate to approximate the composite VAR2CSA sequence of ID1-ID2a_M1010 was not available.

### Monkeys

Female *Aotus nancymaae* monkeys aged 2 to 13 years old were obtained from Keeling center (University of Texas MD Anderson Cancer Center, Michale E. Keeling Center for Comparative Medicine and Research). Animals were housed in stainless steel 6·0 square-foot cages with PVC nesting boxes and wood perches, with a 12:12 dark/light hour photoperiod cycle and room temperature at 24°C. Animals were pair-housed male/female. Standard husbandry procedures included feeding Teklad New World Primate Diet, Zupreem Primate Diet Canned, diet supplements, and water ad-libitum. The monkeys were housed and cared for according to the “Guide for the Care and Use of Laboratory Animals” (ILAR, 2011), Animal Welfare Act and Animal Welfare Regulations (AWA, 2013; AWR, 2013). All animals were enrolled under the Institutional Animal Care and Use Committee-approved malaria candidate vaccine study. The Laboratory of Malaria Immunology and Vaccinology (LMIV), as part of the Public Health Service, Department of Health and Human Services, NIH Intramural Research Program, is accredited by the Association for the Assessment and Accreditation of Laboratory Animal Care, and holds a PHS Assurance on file with the National Institute of Health, Office of Laboratory Animal Welfare as required by the US Public Health Service Policy on Human Care and Use of Laboratory Animals.

### Vaccines and formulations

Vaccines were based on the N-terminal constructs of VAR2CSA from three variants of *P. falciparum* (**Fig. 1A**). The PAMVAC vaccine candidate was generated from the ID1-DBL2-ID2a subunits of the FCR3 allele of VAR2CSA (17). The PRIMVAC vaccine was designed to encompass the DBL1-DBL2 fragment of the 3D7 allele of VAR2CSA (12). A composite VAR2CSA sequence derived from Illumina sequence pileups from the polyclonal maternal isolate PfM1010 was used to generate a vaccine protein from the ID1-ID2a fragment. Briefly, baculovirus (Bv) expression clone and high titer virus stock was produced by Protein Expression Laboratory (NCI-Leidos, Frederick, MD) using a synthetic codon optimized gene encoding ID1-ID2a_M1010 (GenBank Accession number KU665625). Recombinant ID1-ID2a_M1010 has a non-native amino-terminal alanine. All five putative N-linked glycosylation sites were mutated from NxT/S to NxA at positions T440, S507, S518, T694 and T732 relative to VAR2CSA_FCR3_. An in-frame His-tag was included on the carboxyl terminal end to facilitate purification. Bench scale production was done in 10 liters bioreactor glass connected to Biostat BDCU controlling system (Sartorius-Stedim, Bohemia, NY). The cells grew in SFX-insect media (Hyclone) to a concentration of 2 x 10^6^ per ml and were infected with 3 MOI of the recombinant virus. The process was conducted at 27 °C and the dissolved oxygen level was kept at 30%. After 48h hours, the culture was harvested, and the supernatants were concentrated and dialyzed into PBS pH 7.4 prior to purification on a Nickel-Sepharose FF column followed by size exclusion chromatography on a Superdex 200 column. The purity and integrity of ID1-ID2a_M1010 was evaluated by SDS-PAGE as well as by reversed-phase HPLC, and their identity confirmed by N-terminal sequence analysis using Edman degradation. Additionally, the theoretical masses were verified by electrospray ionization mass spectrometry (ESI-MS) (Research Technologies Branch, NIH, Rockville, MD USA). Solution mass and aggregation profile was assessed by analytical size exclusion chromatography with in-line multiangle light scattering (SEC-MALS) as presented in **Fig. S1.**

In addition to these three VAR2CSA-based vaccine candidates, the *Pichia pastoris*-expressed recombinant Pfs25 (a transmission blocking vaccine candidate) (31, 32) was used as a non-VAR2CSA-based control vaccine.

The capsid-based cVLP was designed by genetically fusing a small tag to the PAMVAC antigen, produced in a stable *Drosophila* S2 cell line and coupling to the AP205 based capsid particle displaying multiple catcher domains enabling isopeptide bond formation as previously described (28, 29). Chemically crosslinked conjugate of ID1-ID2a_M1010 with EPA was synthesized as described (30). Briefly, the nanoparticle was prepared by thioether chemistry where sulfhydryl modified ID1-ID2a_M1010 (molecular weight: 68,601 Da) was conjugated with maleimide modified EPA. Sulfhydryl modification of ID1-ID2a_M1010 was obtained by reacting 2·2 mg of ID1-ID2a_M1010 in 1·1 ml of pH 7·2 PBSE/0·5M Guanidinium with 0·22 mg of N-succinimidyl-S-acetyl-thioacetate (SATA) dissolved in DMSO. The mixed reaction was then dialyzed, buffer exchanged into pH6·5 PBSE/0·5M Guanidium buffer using CFD10 centrifugal filter and treated with a deacetylation buffer containing 0·5M hydroxyl amine in pH 7·2 PBSE before another buffer exchanged into pH 6·5 PBSE/0·5M Guanidium buffer. Resulting thiol modified ID1-ID2a_M1010 (1·9 mg) was found to have an average of 4·43 thiols per molecule by DTDP assay. Maleimide modified EPA was generated with 2·85 mg of EPA (molecular weight 66,983 Da) in 0·98 ml pH7·5 PBSE reacting with 0·52 mg of N-ε-malemidocaproyl-oxysuccinimide ester (EMCS) dissolved in DMSO for 90 minutes at 22°C with stirring. Reaction mixture was dialyzed, and buffer exchanged into pH 7·2 PBSE/0·5 Guanidium. The resulting protein mixture (2·74 mg) had a maleimide modification of 6·2 per molecule by reverse DTDP assay. Nanoparticle ID1-ID2a_M1010EPA was prepared by reacting maleimide-modified EPA with thiol-modified ID1-ID2a_M1010. The reaction mixture was stirred at 22°C for one hour. Unreacted maleimide was quenched by adding 21 µl of cystine hydrochloride (0·8 mg/ml) in pH 6·5 PBSE. Conjugate was purified by size exclusion chromatography and analyzed using SEC-MALS for average molecular weight (average molecular weight 1,215 kDa with average diameter by DLS 48 nm) and by amino acid analysis for conjugate composition (56·6% w/w ID1-ID2a_M1010) as described in (30).

Vaccines used in the primary series of vaccination and the booster dose were formulated on Alhydrogel® platform. Details of the different formulations are presented in **Table 1**. Primary vaccine formulations contain the stated doses (antigen content in a volume of 0.5 mL) (**Table 1**A). Nanoparticles PAMVAC_cVLP_ at 12·5 µg/0·5 mL (amount of target antigen) and ID1-ID2a_M1010_EPA_ at 50 µg/0·5 mL as well as the corresponding monomer antigens (PAMVAC and ID1-ID2a_M1010) at 50 µg/0·5 mL doses on Alhydrogel® were injected to the PMV-immunized monkeys (**Table 1**B). Vaccines for Group (1A & 2A) as well as Group (1B & 2B) were formulated by mixing 2X stocks of antigen in their appropriate buffers with 2X stocks of Alhydrogel® in water for injection (WFI) on the days of immunization. Vaccines for Groups 3A and 3B were adsorbed on aluminum at least 24 hours prior to immunization in a final aluminum concentration of 0·404 mg/0·5 mL (=0·808 mg/mL).

**Table 1.** Vaccine treatment groups and formulations. a) Formulation of the vaccines for the primary series of vaccination. b) Booster dose vaccine formulation with antigen content per volume of 0·5 mL indicated.

| **Groups** | **Number of monkeys** | **Vaccine Formulation** | ***Pf* variant** | **Dose (µg/0.5 mL)** | **Alhydrogel (Aluminum content)** |
| --- | --- | --- | --- | --- | --- |
| **1** | **9** | **PAMVAC** | **FCR3** | **50** | **0.85mg** |
| **2** | **9** | **PRIMVAC** | **3D7** | **50** | **0.85mg** |
| **3** | **9** | **ID1-ID2a-M1010** | **M1010** | **50** | **0.404 mg** |
| **4** | **13** | **Pfs25** | **3D7** | **50** | **0.404 mg** |

**b**
| **Groups** | **Formulation Vac 1-3** | **Group # for Dose 4** | **Formulation Dose 4** | **Number of monkeys** | **Antigen Dose (µg/0.5 mL)** | **Aluminum Dose (µg/0.5 mL)** |
| --- | --- | --- | --- | --- | --- | --- |
| **1** | **PAMVAC** | **1A** | PAMVAC Monomer | 3 | 50 | 850 |
|  |  | **1B** | PAMVAC CLP | 4 | 12.5 | 850 |
| **2** | **PRIMVAC (3D7)** | **2A** | PAMVAC Monomer | 3 | 50 | 850 |
|  |  | **2B** | PAMVAC CLP | 3 | 12.5 | 850 |
| **3** | **ID1-ID2a_M1010** | **3A** | ID1-ID2a_M1010 | 2 | 50 | 404 |
|  |  | **3B** | ID1-ID2a_M1010-EPA | 3 | 50 | 404 |

### ELISA titers of PMV-induced antibody in *Aotus* monkeys

Antibody titers induced by PMV candidates as well as the control antigen (Pfs25) were measured by ELISA as previously described (33, 34). Briefly, flat-bottom 96-well ELISA plates (Immunolon 4; Dynex Technology Inc., Chantilly, VA) were coated with 100 ng per well with PAMVAC, PRIMVAC, ID1-ID2a_M1010 and Pfs25 antigens diluted in a carbonate buffer (15mM sodium carbonate and 35mM sodium bicarbonate). Plates were incubated at 4°C overnight and blocked with 200µL/well of the blocking buffer (5% skim milk powder in 1X Tris buffered saline) for 2h at room temperature (RT). *Aotus* serum samples were diluted at 1:500 in blocking buffer, added to duplicated antigen-coated wells (100 L/well) and incubated for 2h at RT. Plates were washed and incubated with 1:3000 dilution of the alkaline phosphatase conjugated anti-Human IgG (H + L) secondary antibody (KPL Inc., Gaithersburg, MD) for 2h at RT. After washing the plates, the substrate (0·1 mg/well of p-nitrophenyl phosphate, Sigma 104 substrate; Sigma) was added to the wells for 20 min incubation in dark at RT, and the absorbance was read at 405nm using a Spectramax 340PC microplate reader (Molecular Devices Co., Sunnyvale, CA).

### Flow cytometry measurement of antiserum reactivity to infected erythrocyte surface

The ability of sera from vaccinated *Aotus* to bind the native antigen expressed on the surface of IE was assessed by flow cytometry. Enriched, mature trophozoite/schizont stages of IE were resuspended in the running buffer (2% of fetal bovine serum in 1X PBS) and 100 µl of the cell suspension was dispensed in each well (4 x 10^5^ cells/well). Cells were incubated with 1:20 dilution of the *Aotus* plasma samples for 30 minutes at 4°C. After unbound antibodies were washed, IE were labeled with 0·1% Sybr Green (Life Technologies) and IgG-bound IE were stained with phycoerythrine-conjugated goat (Fc γ-specific) anti-human IgG (eBioscience) for 30 minutes at 4°C, then washed. Data were acquired by LSRII (BD Bioscience, San Jose, CA) and analysed in FlowJo 10 software (Tree Star Inc.). For each test sample, the ratio median fluorescence intensity (rMFI) was obtained by normalizing the MFI value of the samples with the MFI of the pool of pre-bleed (D0) plasma samples. For qualitative analysis, a ratio greater than 1·2 was considered positive for parasite surface staining (16). Pooled plasma from multigravidae and a VAR2CSA-specific human monoclonal antibody were included in each assay to confirm surface expression of VAR2CSA by the isolates.

### Binding inhibition activity

The CSA binding inhibition capacity of sera from vaccinated *Aotus* was assessed by a static binding inhibition assay on immobilized CSA receptor. Briefly, spots of Decorin (Sigma) at 2 µg/mL in 1X PBS were coated in a 100×15 mm Petri dish (Falcon 351029) and incubated overnight at 4°C in a humid chamber. Spots were then blocked with 3% BSA-1X PBS at 37°C for 30 min. Enriched, mature trophozoite/schizont stages of CS2 and NF54 IE were adjusted to 20% parasite density at 0.5% hematocrit. IE were blocked in 3% BSA-RPMI for 30 min at RT and incubated with *Aotus* plasma at 1:5 dilution for 30 min at 37°C. IE cells were then added to duplicated spots and allowed to settle for 30 min at RT. Unbound IE were washed and adherent IE were immediately fixed with 1·5% glutaraldehyde for 10 min, stained with 5% Giemsa for 5 min and quantified by microscopy. The percentage of inhibition was determined relative to the well with the D0 pre-bleed pool. For a given test sample, the percentage of inhibition was calculated as follows: %inhibition■=■100■−■(Bound-IE_testsample_/ Bound-IE_D0-pool_)■×■100.

### Statistical Analysis

A computer-generated allocation table was used to randomize *Aotus* monkeys into each vaccination group by an independent party. Analysis of the immunological data was executed using GraphPad Prism 8. The activity of antibodies raised against each antigen measured by ELISA, Flow and Binding inhibition assays, were compared using the Mann-Whitney test. Wilcoxon matched-pairs signed rank test was used to evaluate cross-reactivity of PMV-induced antibodies to heterologous antigens and isolates, as well as antibody-boosting activity by CS2 inoculation during pregnancy and PMV-conjugated antigens. Spearman’s correlation coefficient (rho) was used to analyze the relationship between ELISA antibody titers and their ability to recognize the native VAR2CSA and the blockade of the parasite binding to CSA. P values <0·05 were considered significant.

## Results

### Local and Systemic Reactogenicity

Overall, vaccines were well-tolerated by the *Aotus* monkeys. No death occurred during the vaccination period. Fifteen out of 40 monkeys showed some muscle firmness at the injection site after vaccination with no edema, redness or warmth. Hardened areas of the muscle were measured using a caliper to determine the time at which the reaction dissipated from the animals. This muscle induration was observed across all vaccination groups, including the Pfs25 control group, and resolved within 7 days post-vaccination. No sign of systemic reaction or change in behavior was observed. Detailed description of clinical observations post-immunization is presented as **Data File S1.**

### PMV-induced antibodies in *Aotus* exhibit strong homologous activity

All 40 monkeys completed vaccination and provided sera two weeks after the last vaccination (D70). Serum samples from vaccinated animals were tested by ELISA for vaccine-specific antibody reactivity and all antisera showed IgG titers against their corresponding antigen (**Fig. 2A**). Antibodies from Pfs25-immunized monkeys showed no cross-reactivity to the VAR2CSA-based antigens. Similarly, none of the PMV-induced antibodies reacted to Pfs25 antigen (**Fig. 2A**). All PMV antisera showed significantly higher cross-reactivity to PMV antigens compared to Pfs25 antisera (Mann Whitney test, P < 0·001 for all comparisons). Overall, PMV candidates induced a high level of vaccine-specific IgG to homologous antigen with some level of cross-reactivity to heterologous PMV antigens (**Fig. 2A** and **Fig. S1**). PAMVAC-induced antibodies showed a similar level of heterologous reactivity to PRIMVAC and ID1-ID2a_M1010 antigens (**Fig. S2**), whereas antibodies from PRIMVAC and ID1-ID2a_M1010 immunizations respectively had higher heterologous reactivity to ID1-ID2a_M1010 and PRIMVAC antigens than to PAMVAC (Wilcoxon matched-pairs signed rank test, P = 0·02 for PRIMVAC-IgG and P = 0·004 for ID1-ID2a_M1010-IgG).

**Figure 2.**
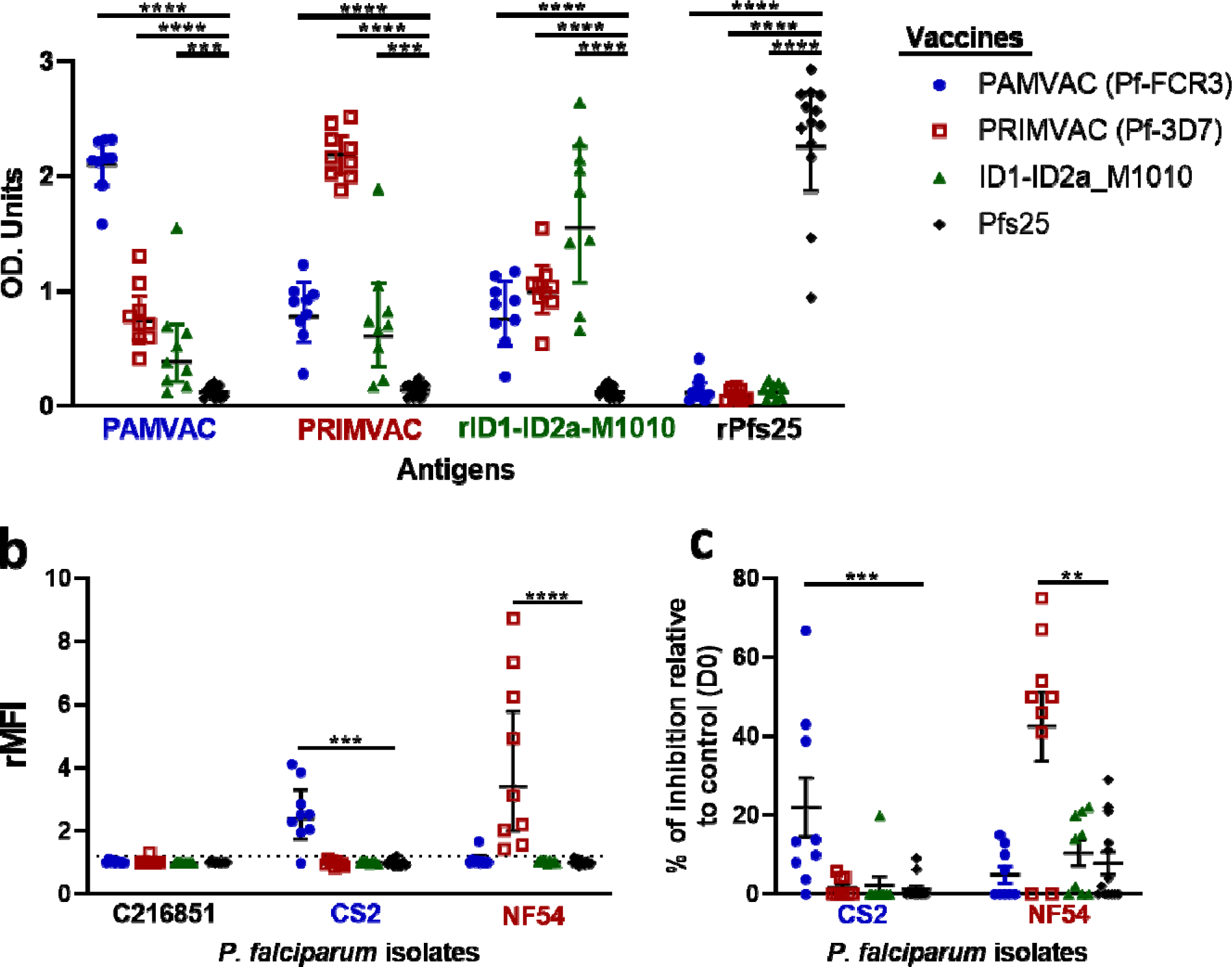
Activities of vaccine-induced antibodies from *Aotus* following immunization. a) The reactivity of vaccine-induced IgG two weeks after the last vaccination dose was assessed against PMV candidates PAMVAC (Pf-FCR3) and PRIMVAC (Pf-3D7) as well as recombinant antigens rID1-ID2a-M1010 and rPfs25. Functional characterization of antibodies was evaluated for b) the ability of IgG to bind antigens expressed on the surface of the IEs by Flow cytometry analysis, and for c) CSA-binding inhibition activity against different *P. falciparum* isolates. For each group of vaccinated monkeys, the geometric mean and 95% CI of the antibody activity measured by ELISA and Flow cytometry are shown. For BIA, the mean and standard error of the mean (SEM) are shown.

CS2 and NF54 isolates were used to assess the reactivity of these antibodies in plasma to native VAR2CSA. For each test sample, the ratio median fluorescence intensity (rMFI) was obtained by normalizing the MFI value of the samples at D70 with the MFI of the pool of pre-bleed (D0) plasma samples. Functional characterization of PMV-induced IgG demonstrated strong ability to recognize the native VAR2CSA expressed by the homologous parasite strain (mean rMFI = 2·4 for PAMVAC group against CS2 and mean rMFI = 3·4 for PRIMVAC group against NF54) (**Fig. 2B**). Unlike the cross-reactivity of PMV-induced IgG to different VAR2CSA-based antigens measured by ELISA, no heterologous reactivity of antibodies binding to VAR2CSA-expressing CS2- and NF54-IE was observed in flow cytometry analysis. No IE surface-labelling of the child isolate (C216851) with CD36-binding phenotype was observed in plasma from the immunized monkeys, and IgG from Pfs25-immunized monkeys showed no reactivity to all three isolates. These findings revealed a weak or absent capacity of PMV-induced antibodies from *Aotus* to recognize epitopes shared by different native antigens.

This pattern of plasma reactivity to native surface antigen corresponded to limited heterologous binding inhibition activity against CS2 and NF54 isolates (**Fig. 2C**). Plasma from PAMVAC-vaccinated monkeys at 1:5 dilution significantly inhibited CS2 binding to CSA (mean % inhibition = 21·9) compared to Pfs25 (mean inhibition = 1·2 %; Mann Whitney test, P = 0·0005), PRIMVAC (mean inhibition = 1·6 %; Mann Whitney test, P = 0·003) and ID1-ID2a_M1010 (mean inhibition = 2·2 %; Mann Whitney test, P = 0·002) vaccine groups. Conversely, PRIMVAC-induced antibodies significantly blocked NF54 binding to CSA (mean inhibition = 42·6 %) compared to Pfs25 (mean inhibition = 7·8 %; Mann Whitney test, P = 0·007), PAMVAC (mean % inhibition = 4·9; Mann Whitney test, P = 0·01) and ID1-ID2a_M1010 (mean % inhibition = 10·4; Mann Whitney test, P = 0·02) induced antibodies.

The relationship between PAMVAC- and PRIMVAC-induced antibody levels and functional activity against CS2 and NF54 isolates was analyzed. Overall, in the PAMVAC vaccine group, the level of antibodies binding to PAMVAC antigens measured by ELISA did not correlate with the level of IE-surface recognition and binding inhibition activity against the homologous parasite (CS2) (**Fig. 3**). In the PRIMVAC group, the level of antibody binding to PRIMVAC antigen strongly correlated with that of IE-surface recognition of NF54 (rho = 0·92; P = 0·001), but not the binding inhibition activity (**Fig. 3**). In addition, the levels of surface reactivity and binding inhibition activity to NF54 were significantly correlated for PRIMVAC-induced antibodies (rho = 0·74; P = 0·03, **Fig. 3**).

**Figure 3.**
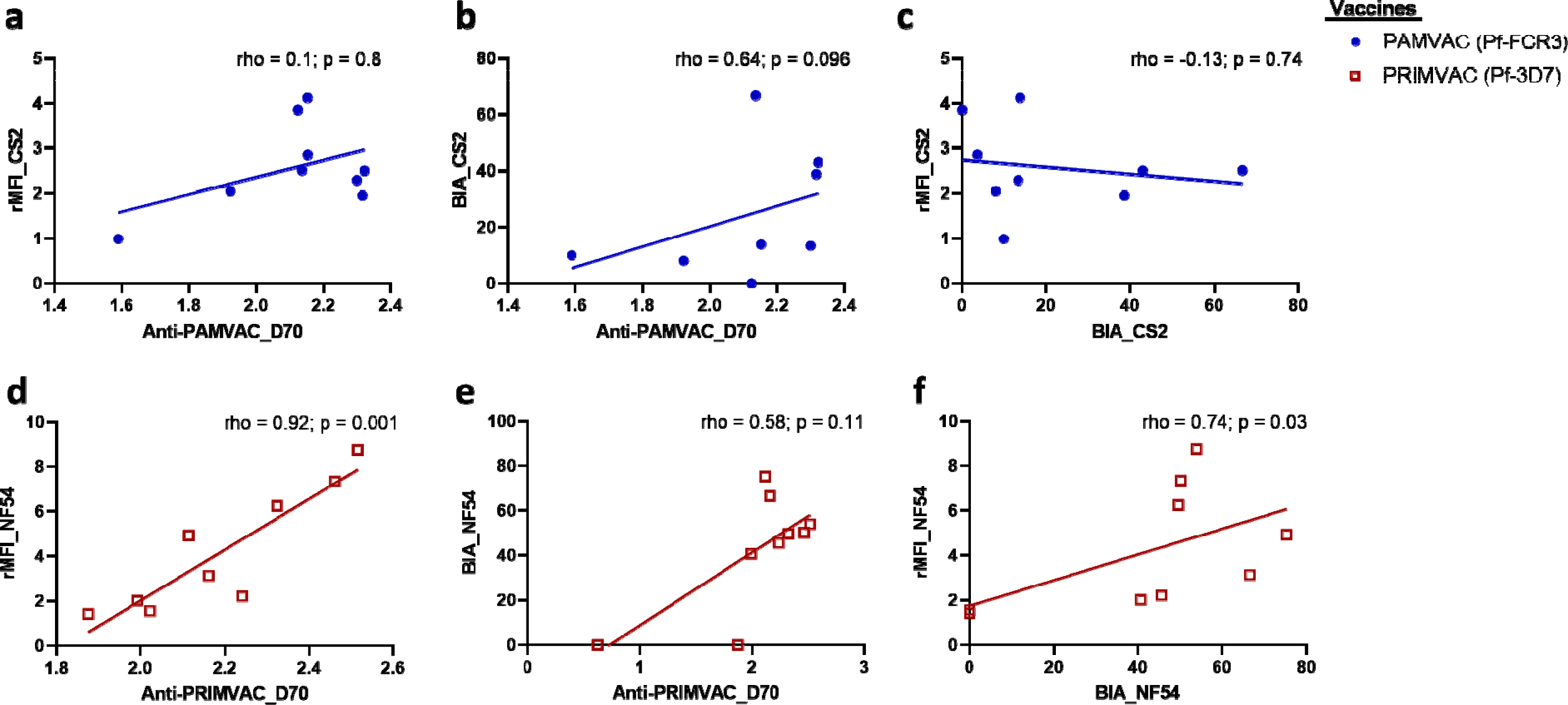
Correlations between ELISA titers of PMV-induced antibodies and functional activity at D70 post-vaccination. Spearman (rho) coefficients and P values are reported to describe the relationship between ELISA titers of PAMVAC antibodies in *Aotus* and surface reactivity (a) or binding inhibition activity (b) against CS2 parasite. The correlation between surface reactivity and BIA of PAMVAC antibodies is also shown (c). The same analyses were performed for PRIMVAC-induced antibodies in *Aotus* and presented in panels d, e and f. Data for animals immunized with PAMVAC (blue circle) and PRIMVAC (open red square) are shown.

### A single acute malaria infection during pregnancy does not boost the activity of PMV-induced antibodies in *Aotus*

The *Aotus* PM model offers a unique opportunity to investigate the impact of *P. falciparum* infection during pregnancy on PMV-specific antibodies. Twenty-three immunized animals became pregnant and were infected with CS2 parasite. For this acute infection, a week-long infection was allowed before C-section was performed.

Antibody reactivity and function were assessed prior to infection and 4 weeks post-delivery. Because pregnancy occurs sporadically in primates, the timing between the D70 post-vaccination serum sample and the day of CS2 inoculation during pregnancy varied by monkey and ranged from 6 to 584 days; however, there was no statistically significant difference between the groups in the median time to CS2 inoculation (**Fig. S3**). The time interval between the day of CS2 inoculation and 4 weeks postpartum was 35 days for all animals. No response was observed in Pfs25-vaccinated animals (measured by ELISA on PMV antigens, flow cytometry and binding inhibition assays on CS2 and NF54) when comparing D70 post-vaccination to 4 weeks postpartum measurements (**Fig. 4**), as might be expected because Pfs25 is not expressed by blood-stage parasites.

**Figure 4.**
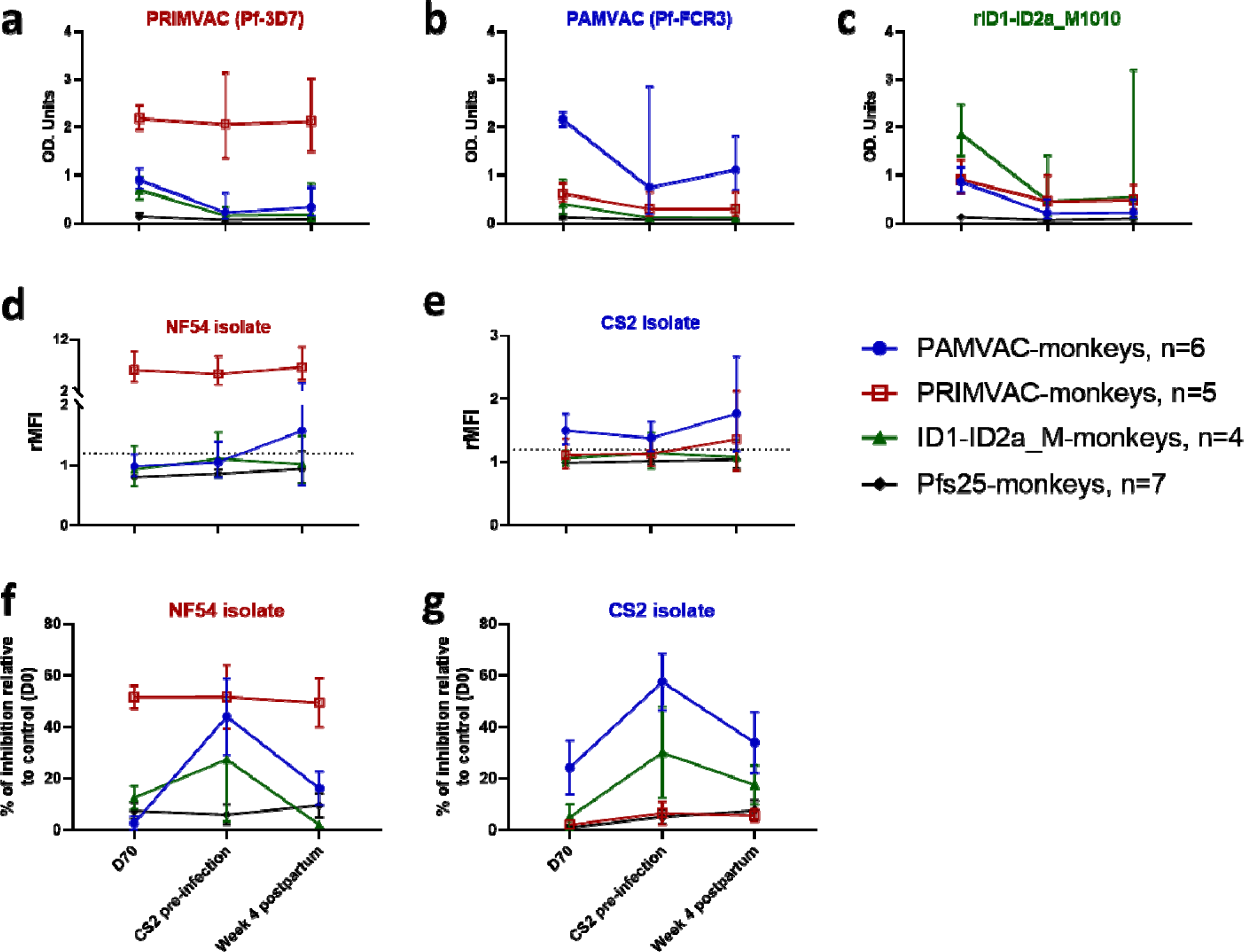
Dynamics of VAR2CSA antibodies following a CS2 inoculation during pregnancy. ELISA titers 2 weeks after the last dose in the primary vaccine series compared to those measured at the time of CS2 parasite inoculation and 4 weeks postpartum, for each of the vaccine groups (panels a, b, and c). Flow cytometry for IE surface reactivity (panels d and e), and BIA for parasite blocking activity (panels f and g). For each group of vaccinated monkeys, the geometric mean and 95% CI of the antibody activity measured by ELISA and Flow cytometry are shown. For BIA, the mean and standard error of the mean (SEM) are shown.

Strikingly, PRIMVAC-induced antibody levels had not declined over the months or years between immunization up to the experimental pregnancy malaria episode (**Fig. 4A**); vaccine responses induced by other PMV antigens had declined in the interregnum (**Fig. 4B, 4C**). Following infection during pregnancy, no significant increase above the pre-infection level was observed by IgG ELISA for any PMV group to its corresponding PMV antigen (**Fig. 4A-C**); a slight non-significant increase in ELISA IgG reactivity of PAMVAC antisera to PAMVAC antigen may have occurred. Similarly, the level of reactivity to NF54 and CS2 IE surface recognition in sera from PRIMVAC- and PAMVAC-immunized monkeys, respectively, did not increase following CS2 infection during pregnancy (Wilcoxon matched-pairs signed rank test, P = 0·06 for PRIMVAC group against NF54 and P = 0·3 for PAMVAC group against CS2) (**Fig. 4D and 4E**).

CSA-binding inhibition activity of antibodies from PMV-vaccinated groups against both homologous and heterologous parasites showed a different trend versus other assays: the level of inhibition from D70 post-vaccination to the day of CS2 inoculation increased (PAMVAC group; Wilcoxon matched-pairs signed rank test, P = 0·03) or was unchanged (**Fig. 4F and 4G**). Unexpectedly, a non-statistically significant decline of this activity was observed at 4 weeks postpartum following CS2 inoculation during pregnancy. Overall, there was no significant change in the activity of PMV-induced antibodies between the D70 post-vaccination and 4 weeks postpartum, a time window ranging from 41-1003 days. Remarkably, all homologous responses (measured by ELISA, Flow and BIA) of PRIMVAC-induced antibodies against PRIMVAC antigen and NF54 parasite appeared stable over time from D70 post-vaccination through 4 weeks postpartum (**Fig. 4**).

The levels of PAMVAC- and PRIMVAC-induced antibodies binding to the homologous PMV antigen did not significantly correlate with the level of IE-surface recognition and binding inhibition activities against homologous parasites at the day of CS2 inoculation (**Fig. S4**) and 4 weeks postpartum (**Fig. S5**).

### PMV can induce a delayed acquisition of cross-inhibitory activity against CS2 and NF54 parasites

To explore strain-transcending activity of PMV-induced antibodies in *Aotus,* we analyzed the cross-inhibitory and cross-recognition activities of the antibodies against both CS2 and NF54 parasites. These activities were assessed on samples collected at D70 post-vaccination, on the day of CS2 inoculation, and 4 weeks postpartum. When defining cross-inhibitory activity as >50% inhibition of both CS2 and NF54 binding to CSA, no serum samples showed such activity at D70 post-vaccination and 4 weeks postpartum (**Fig. 5A and 5C**). Three monkeys (*Aotus* AO12 and AO10 that received PAMVAC and AO13 that received ID1-ID2a_M1010 vaccine, and were respectively inoculated with CS2 8, 346 and 381 days after D70 post-vaccination samples were collected) had developed cross-inhibitory activity against CS2 and NF54 measured just prior to CS2 inoculation (**Fig. 5B**). Curiously, this cross-inhibitory activity waned by 4 weeks postpartum despite exposure to CS2 inoculation (**Fig. 5C**). Of note, some other monkeys maintained their homologous inhibitory activity from D70 post-vaccination through 4 weeks postpartum (*Aotus* AO1 from PAMVAC group; AO4 and AO5 from PRIMVAC group), while others showed newly acquired homologous activity at the time of CS2 inoculation that persisted at 4 weeks postpartum (*Aotus* AO11 and AO8 from PRIMVAC group).

**Figure 5.**
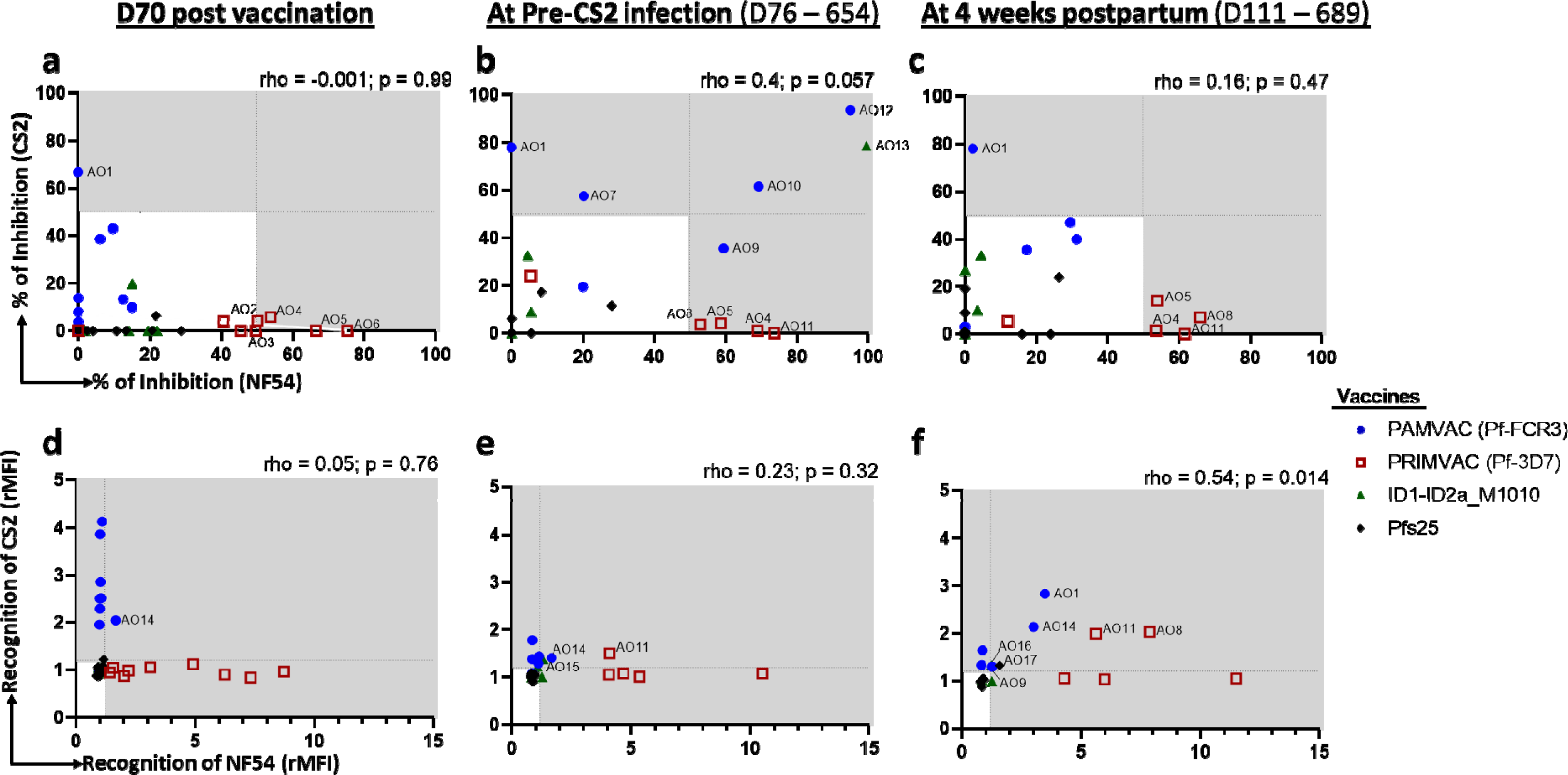
Dynamics of cross-inhibitory and cross-recognition activity of PMV-induced antibodies in *Aotus* monkeys in response to pregnancy malaria. The levels of binding inhibition activity of the PMV-induced antibodies against CS2 and NF54 parasites in samples collected at D70 post-vaccination (panel a), Pre-CS2 inoculation (panel b) and 4 weeks postpartum (panel c) are shown. The dashed lines represent 50% inhibition and the gray colored area in each graph defines labeled samples with inhibitory activity. The recognition levels of surface-expressed VAR2CSA on CS2 and NF54 IE by the PMV-induced antibodies was also analyzed at D70 post-vaccination (panel d), Pre-CS2 inoculation (panel e) and 4 weeks postpartum (panel f), with dashed lines representing rMFI = 1·2 and gray colored area highlighting Flow positives samples. Data for animals immunized with PAMVAC (Pf-FCR3) (blue circle), PRIMVAC (Pf-3D7) (open red square), rID1-ID2a-M1010 (green triangle) and rPfs25 (black diamond) are shown and samples with cross-recognition activity are labeled. Spearman (rho) coefficients and P values are reported for each graph.

Surface recognition of native VAR2CSA expressed on CS2 and NF54 IE followed a pattern different than that observed for the inhibitory activity profile. Limited cross-recognition activity of the PMV-induced antibodies in *Aotus* was detected at D70 post-vaccination, with only one monkey (AO14 from PAMVAC group) exhibiting such activity (**Fig. 5D**). By the time of CS2 inoculation, two additional monkeys (AO11 from PRIMVAC group and AO15 from ID1-ID2a_M1010 group) had acquired the cross-recognition activity (**Fig. 5E**). Following CS2 inoculation, four additional monkeys (AO9 and AO1 from PAMVAC group, AO11 and AO8 from PRIMVAC group, AO17 from Pfs25 group), developed IE surface cross-recognition activity (**Fig. 5F**). Such cross-recognition activity detected on CS2 and NF54 parasites five weeks after exposure to CS2 was significantly correlated (rho = 0·54; P = 0·014, **Fig. 5F**).

### Nanoparticle or monomer PMV similarly boost ELISA reactivity and not function

On study Day 999 (cohort 1) and 853 (cohort 2) (943 and 797 days after completing the primary vaccination series for cohort 1 and 2, respectively), 18 monkeys were immunized with either nanoparticle (PAMVAC_cVLP_ or ID1-ID2a_M1010_EPA_) or monomer forms of PAMVAC or ID1-ID2a_M1010. Before re-immunization (pre-boost), PAMVAC monkeys displayed minimal or modest antibody responses to homologous antigen (**Fig. 6A**) while PRIMVAC monkeys maintained substantial levels of antibody to homologous antigen (**Fig. 6B**). Fourteen days after administration, serum IgG binding to PMV antigens by ELISA substantially increased in all groups, with highest reactivity to the antigen used for the primary vaccine series and high cross-reactivity to other antigens (**Fig. 6A, 6B and 6C**). PMV-induced antibody levels following the primary immunization series (Day 70) had waned over time but the late booster dose increased levels beyond those achieved by primary series (**Fig. S6**). Homologous vaccine boosted reactivity for PAMVAC- and ID1-ID2a_M1010-induced antibodies, and the heterologous vaccine boosted reactivity of PRIMVAC-induced antibodies as well. Indeed, the PAMVAC boost of the PRIMVAC-vaccinated group substantially increased the levels of antibodies against all three PMV. Of note, the monomer and the conjugated vaccines similarly increased ELISA reactivity at D+14 and D+56 post-booster dose (Mann Whitney test, P > 0·05 for all comparisons).

**Figure 6.**
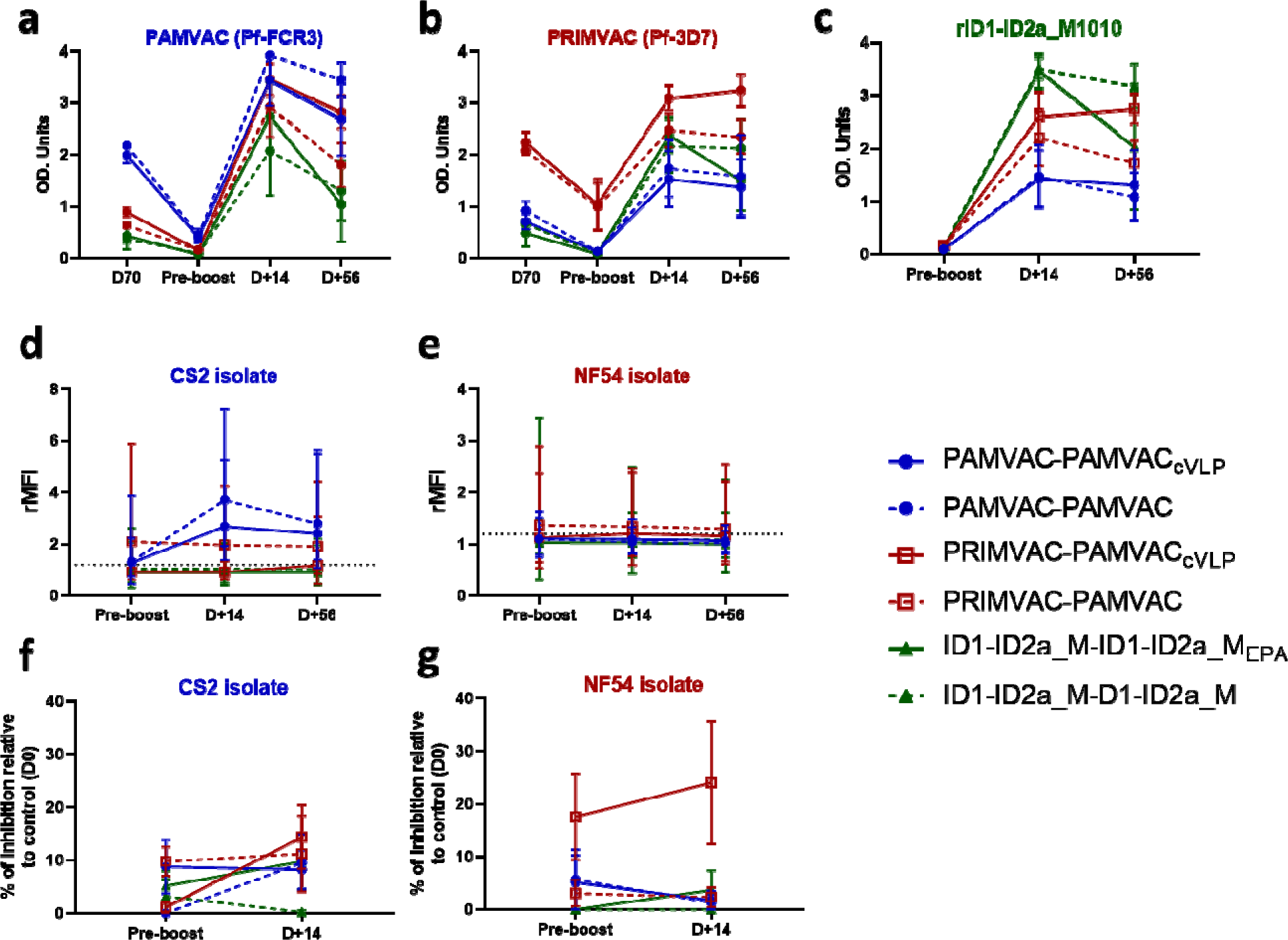
Activities of PMV-induced antibodies in *Aotus* monkeys following re-vaccination with monomer or nanoparticle immunogens. ELISA titer of IgG against a) PAMVAC (Pf-FCR3), b) PRIMVAC (Pf-3D7) and c) ID1-ID2a-M1010 antigens were assessed in monkeys that received the conjugated antigens (solid-line, PAMVAC_cVLP_ or ID1-ID2a-M1010-EPA) and those who received the monomer antigens (dashed-line). Samples collected before vaccination (Pre-bleed) and those collected at D14 and D56 post-vaccination were analyzed. Plasma samples collected at the same timepoints were also analyzed for their surface reactivity to d) CS2 and e) NF54 IEs, as well as their ability to block those parasites binding to CSA (f and g). For each group of vaccinated monkeys, the geometric mean and 95% CI of the antibody activity measured by Flow cytometry are shown. For ELISA and BIA, the mean and standard error of the mean (SEM) are shown.

Antibody reactivity to native VAR2CSA expressed by CS2 and NF54 isolates showed no change in groups initially vaccinated with PRIMVAC and ID1-ID2a_M1010 for both isolates (**Fig. 6D and 6E**), while PAMVAC-vaccinated monkeys showed an increase in reactivity to homologous CS2-IE (Mann Whitney test, P = 0·05 for PAMVAC_cVLP_ boost group and P > 0·05 for PAMVAC boost group, both at D+14 and D+56 post-booster dose). The nanoparticles did not significantly increase the plasma binding inhibition activity against CS2 and NF54 at D+14 for any group (**Fig. 6F and 6G**), albeit serum activity of PRIMVAC-immunized animals was higher against both parasites after the booster dose, but still modest (less than 40%).

## Discussion

Ideally, PMV will be administered to women before their first pregnancy in order to induce protective antibodies similar to those in PM-resistant multigravid women and will be boosted on exposure to infection during pregnancy (6, 35, 36). Several lines of evidence support VAR2CSA as the leading PMV candidate to protect pregnant women from PM-related adverse pregnancy outcomes (9, 11, 13, 15, 35, 37-41). Recently published first-in-human trials of VAR2CSA-based vaccines established that adjuvanted PAMVAC and PRIMVAC are safe and immunogenic in malaria-naïve and malaria-exposed women (21, 22). Although both PAMVAC and PRIMVAC vaccines adjuvanted with Alhydrogel® induced high human antibody titers, functional activity was generally modest and limited to homologous variants of the vaccines (21, 22, 42). The present study demonstrates the ability of the *Aotus nancymaae* model to predict the human immune response to VAR2CSA-based vaccination.

A NHP model susceptible to *P. falciparum* placental infection will be useful to study PM pathogenesis and evaluate vaccine activity (reviewed in (43)). The establishment of an *Aotus* model that displays key features of PM pathogenesis, such as placental sequestration and the acquisition of functional antibodies following exposure to placenta-binding parasites (Sharma et al., submitted), represents a major advance to support PMV development. The current data demonstrate that Alhydrogel® formulated PAMVAC, PRIMVAC and the ID1-ID2a construct of isolate M1010 were safe in *Aotus* monkeys and induced high levels of anti-VAR2CSA antibodies against the corresponding vaccine antigen, with some cross-reactivity against heterologous PMV antigens by ELISA. These findings are consistent with previous reports from PAMVAC and PRIMVAC preclinical studies in small animals (12, 17, 20) and human clinical trials (21, 22), highlighting the safety and immunogenicity of VAR2CSA-based sub-unit vaccines in both soluble and particle-based forms. Cross-reactivity of PMV-induced antibodies against the vaccine antigens indicates that the induction of antibodies targeting epitopes shared between different recombinant PMV variants is not dependent on the host species.

However, cross-reactivity by ELISA did not correspond to broadly functional antibodies. Contrary to rodent studies of many VAR2CSA candidates including PRIMVAC and PAMVAC (11-19), VAR2CSA antisera from *Aotus* poorly reacted to native VAR2CSA expressed on the surface of heterologous parasite strains while reacting strongly to homologous parasites.

Similarly, *Aotus* antisera blocked CSA binding of homologous parasite strains but not heterologous parasites with up to 66·7 % homologous inhibition activity in the PAMVAC group and 75·0% in the PRIMVAC group. This lack of cross-recognition and cross-inhibition activity substantiates the functional pattern of antibodies induced by both vaccines using the same dosage and adjuvant in women (21, 22). Although a higher vaccine dose and a more potent adjuvant might yield higher activity, in humans 100 µg of PRIMVAC adjuvanted with Alhydrogel® or GLA-SE yielded limited cross-recognition against the FCR3-CSA and 7G8-CSA VAR2CSA-expressing parasites (22). Overall, these observations clearly demonstrate that antibody activity induced by PMV antigens in humans aligns better to that induced in *Aotus* than in small animal models. The overlapping profile of antigens recognized by antibodies from human and *Aotus* after malaria infection (25) could explain the high similarity in VAR2CSA-specific antibody response between these two hosts.

Antibodies to VAR2CSA are thought to be boosted through repeated exposure to placenta-binding parasites over successive pregnancies (35, 44, 45). By exploring whether a malaria infection in pregnant *Aotus* monkeys immunized with PMV antigens will enhance the functional properties of the induced VAR2CSA-specific antibodies, we showed that a single week-long infection with CS2 did not significantly boost antibody function at 4 weeks postpartum. Similarly, CS2 infection in pregnant monkeys vaccinated with Pfs25 did not induce antibodies to recombinant VAR2CSA or CSA-binding parasites. Since the PfCS2 inoculi used for infection were predominantly binding to CD36 receptor, these observations suggest that few IEs had CSA-binding phenotype and therefore animals might have experienced a short infection insufficient to mount a significant antibody response to VAR2CSA. As most malaria infections in pregnant women are sub-patent and asymptomatic (46), it is possible that a longer duration of malaria infection may be needed to induce/enhance functional properties of VAR2CSA antibodies. To that end, a chronic model of PM in *Aotus* is currently under development and will be used to investigate whether chronic malaria infection in pregnant monkeys can induce/enhance functional activity of VAR2CSA antibodies.

Although no significant boost of PMV-induced antibodies occurred after CS2 inoculation during pregnancy, the high level of PRIMVAC-induced homologous functional antibodies persisted from D70 to 4 weeks postpartum in *Aotus* (up to 619 Days), reproducing the observed long-term seroconversion in women after the last PRIMVAC immunization (22). This finding indicates that VAR2CSA-specific antibodies induced by vaccination may be durable, and their functional activity could be expected to be sustained across pregnancies. This result is particularly important since the targeted population will be nulligravid women, and first pregnancy could occur long after the vaccination. Naturally acquired human VAR2CSA antibody responses are similarly durable, with antibody half-life estimated to be years in duration (45, 47).

The antibodies naturally acquired by multigravidae have strain-transcending functional activity against placenta-binding IE (6), and VAR2CSA is thought to be the target of these antibodies (reviewed in (48)). Overall, in *Aotus*, the pattern of cross-inhibitory activity (measured by BIA) differed from cross-recognition of native VAR2CSA expressed by CS2 and NF54 (measured by flow cytometry), as many monkeys with IE surface cross-recognition had no cross-inhibitory activity. A number of monkeys acquired cross-recognition activity of antibodies between their post-primary vaccination timepoint (D70) and 4 weeks postpartum, while transient cross-inhibition activity (for 2 monkeys in PAMVAC and 1 monkey in ID1-ID2a_M1010 groups) was detected. Since the durability of VAR2CSA-specific inhibitory antibodies are unclear, future investigations of the dynamics of inhibitory antibody generation in this *Aotus* model will be of interest. Because immunological memory specific for VAR2CSA can be maintained for many years without antigen re-exposure (49), it is conceivable that affinity maturation yielding antibody with higher avidity may play a key role in the inhibitory and strain-transcending functional activities of VAR2CSA-induced antibodies (50, 51).

Efforts to optimize malaria vaccines (including PMV) include use of cVLP and EPA nanoparticle platforms to enhance immunogenicity (21, 26, 52). Here, PAMVAC and ID1-ID2a_M1010 nanoparticles generated in cVLP and EPA platforms, respectively, boosted antibody levels measured by ELISA above those seen after the primary vaccine series. However, it appeared that monomeric antigens boosted antibody responses equally well, suggesting this boosted response is not improved by nanoparticle presentation. Notably, heterologous PAMVAC boost of PRIMVAC-vaccinated *Aotus* monkeys substantially enhanced levels of antibodies against all three PMV antigens. However, the increase in ELISA titers induced by PMV (whether nanoparticle or monomer) did not correspond to an increase in functional activity assessed by flow cytometry and BIA, suggesting qualitative differences in VAR2CSA antibody responses. A head-to-head comparison of the cVLP-based and monomer vaccines for priming versus boosting in future *Aotus* studies will provide a useful indication for how a nanoparticle-based vaccine might behave in humans.

This study had limitations. One practical limitation of the *Aotus* model is the variable and sometimes long-time window between vaccination and onset of pregnancy, unlike rodent models that more reliably reproduce. However, the *Aotus* is similar to what can be expected in humans, in whom timing of pregnancy is also variable. Therefore, this *Aotus* model may require extended time periods to assess vaccine boosting and efficacy to pregnancy infection. However, this may also be a model to understand immunological memory, as is also important in humans. As another limitation, the current study had limited power to assess placental parasite burden: because the *Aotus* practices placentophagy, a C-section must be performed to secure the placental sample. In this study, only few paired peripheral and placental parasitemia were obtained, and this weakened statistical power to determine any significant patterns, although general trend suggests a higher parasite burden in the placenta compared to peripheral blood irrespective of the immunization groups (**Fig. S7**). The *Aotus* model has now been improved to define optimal timing for C-section, which will allow systematic collection of placental samples in future studies. In addition to the model of chronic placental malaria in development, these improvements should strengthen the use of the *Aotus* model as a predictive model for vaccine-induced PM protection in pregnant women. Finally, due to logistical and regulatory requirements for the use of NHP models, accessibility of this *Aotus* model of PM may be limited to centers with the necessary infrastructure. Other animal models with human immunoglobulin loci or wild-type small rodent models can also be explored, in addition to *Aotus*, in ongoing efforts to improve strain-transcending functional activity of PMV-induced antibodies.

In conclusion, we demonstrated that the *Aotus* model is a suitable model to assess immunogenicity of VAR2CSA-derived vaccines, in contrast to small animal models. Current PMV candidates induce mainly homologous and little heterologous functional activity in humans and *Aotus*, suggesting that improvements to the immunogens and/or the adjuvants are needed to enhance protective antibody responses, as are studies that evaluate the potential for natural infection to boost vaccine antibody in women. The *Aotus* model further suggests that a brief PM episode may not boost PMV functional activity, and that PMV nanoparticles are not superior to monomers as immunogens for boosting. Further work with the *Aotus* model should assess placental parasite burden as an endpoint for interventional studies, as well as the ability of chronic placental malaria episodes to boost VAR2CSA functional antibodies. The *Aotus* PM model may be useful to assess second-generation PMVs seeking to increase strain-transcending activity and prioritize these for further clinical development.

## Author contributions

JD, MAN, NKV, SOH, MF, TGT, BG and PED conceived of the study, designed the studies, and interpreted data. LL, MCN, ASalanti, CA, KR, DLN and PVS contributed to study design and data interpretation. JD, LEL, AC, AFC, JPS, SOG, CMJ, AJS, SBC, MSS, EB, JS, BBC, SN, KH, TO, SC, ASharma, HT, BB and PVS performed study experiments. MLT provided medical and surgical care to the animals. JD, MF and PED analyzed data. JD and PED wrote the manuscript with input from all authors and approval for the final version.

## Acknowledgements

This work was supported by the Intramural Research Program of the National Institute of Allergy and Infectious Diseases, National Institutes of Health; the Danish Research Councils, The Eurostars Programme, the European Union in the Seventh program Framework Programme (FP7-HEALTH-2012-INNOVATION; under grant agreement 304815), the Danish Advanced Technology Foundation (under grant number 005-2011-1), the Bill and Melinda Gates Foundation (grant numbers 42387 and OPP1055855 to S. G. R.); the French National Research Agency (ANR-16-CE11-0014-01), grants from Laboratory of Excellence GR-Ex, reference ANR-11-LABX-0051 and the French Parasitology consortium ParaFrap (ANR-11-LABX0024). The labex GR-Ex is funded by the program “Investissements d’avenir” of the French National Research Agency, reference ANR-11-IDEX-0005-02; a grant from the Bundesministerium für Bildung und Forschung (BMBF), Germany through Kreditanstalt für Wiederaufbau (KfW) (Reference No: 202060457) and through funding from Irish Aid, Department of Foreign Affairs and Trade, Ireland.

The authors thank J. Patrick Gorres for editing the manuscript. We are grateful to Sharon Wong-Madden, Jennifer Kwan, Matthew Cowles, Rhea Stevens, Robert Morrison and Jill Neal for their technical assistance and advice, and Fabrice A. Somé, Maria del Mar Castro Noriega and Moussa Niangaly for project management assistance.

## Data Sharing Statement

All data associated with this study are present in the paper or Supplementary Materials and are available from the authors upon reasonable request.

