## Supplemental Material for "*Aotus nancymaae* model predicts human immune response to the placental malaria vaccine candidate VAR2CSA"

**Supplemental Figure S1: Biochemical and biophysical characterizations of recombinant ID1-ID2a_M1010 protein for identity, integrity and purity.**

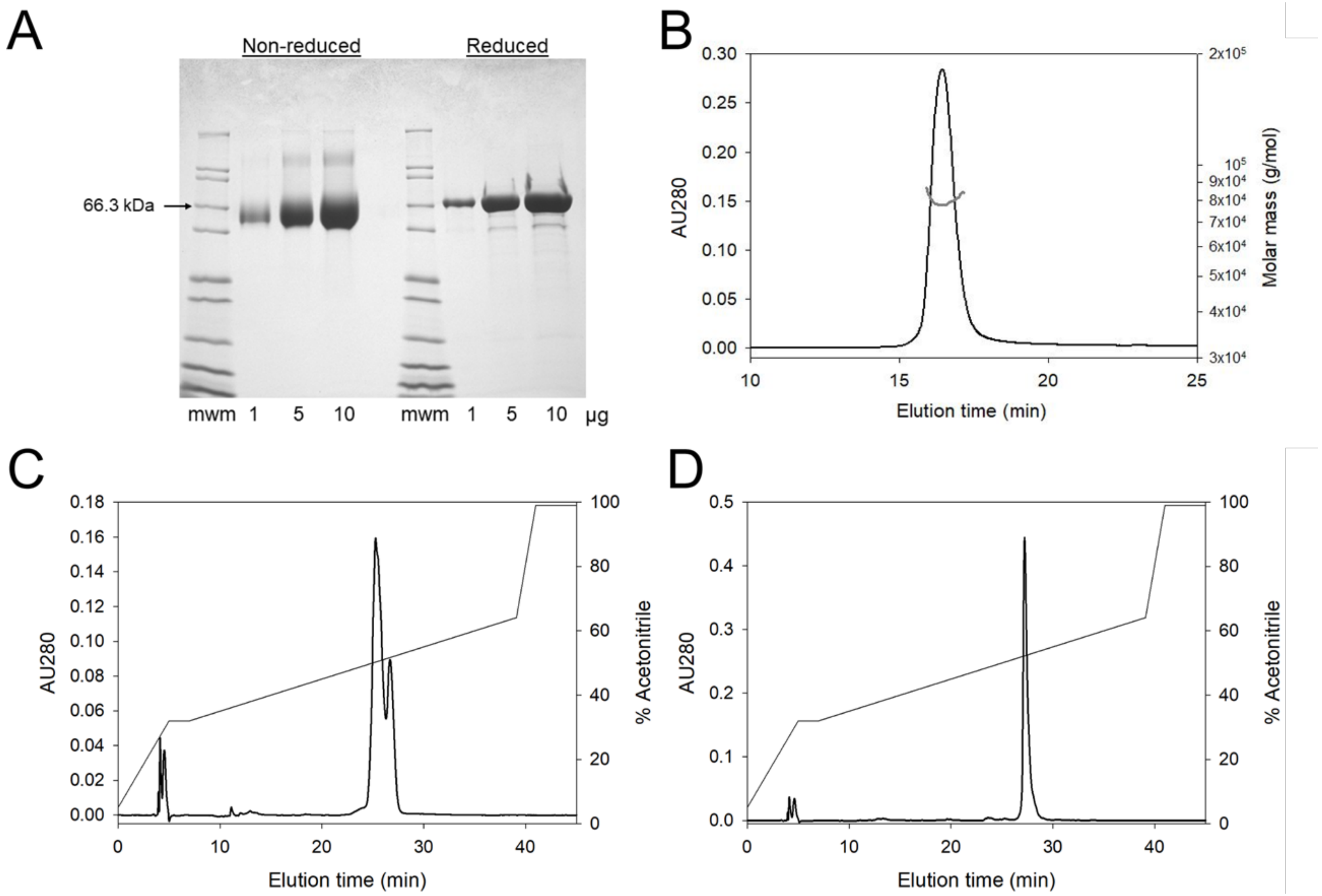

**Supplemental Figure S2: ELISA cross-reactivity of PMV-induced antibodies among VAR2CSA antigens.**

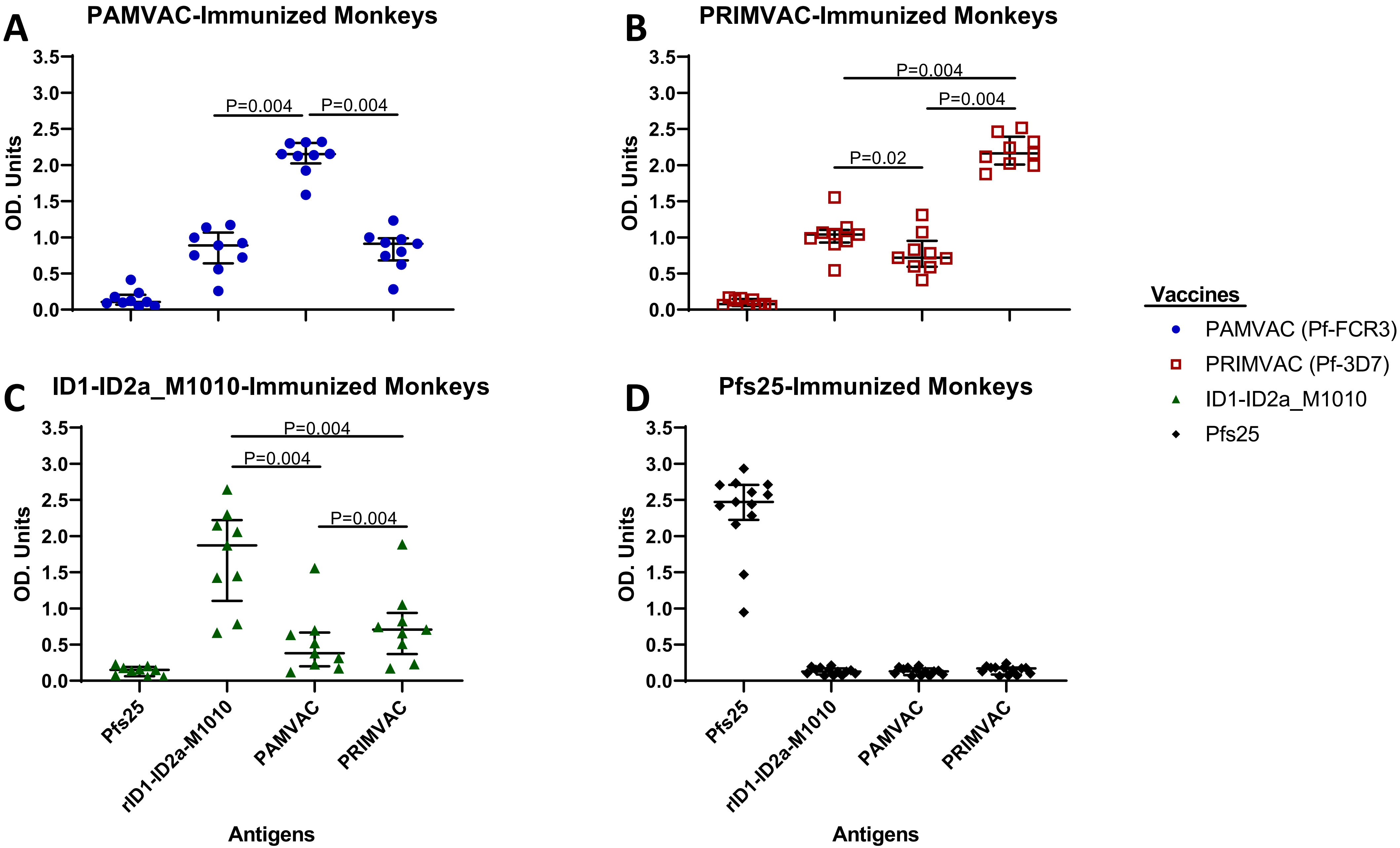

**Supplemental Figure S3: Difference in time from D70 post vaccination 1 to CS2 infection**

A

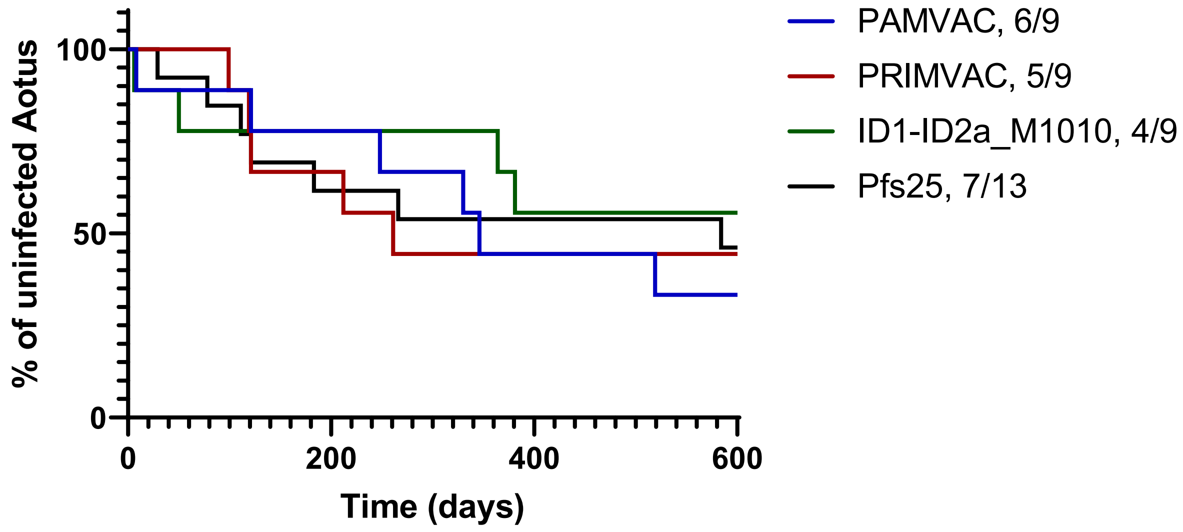


B

| Comparison of vaccine groups (x vs. y) | Medians (x / y) | P-value |
|---|---|---|
| ID1-ID2a_M1010 vs. PAMVAC | 207 / 289 | 0.91 |
| ID1-ID2a_M1010 vs. Pfs25 | 207 / 121 | 0.93 |
| ID1-ID2a_M1010 vs. PRIMVAC | 207 / 121 | 1 |
| PAMVAC vs. Pfs25 | 289 / 121 | 0.47 |
| PAMVAC vs. PRIMVAC | 289 / 121 | 0.27 |
| Pfs25 vs. PRIMVAC | 121 / 121 | 0.93 |

**Supplemental Figure S4: Correlations between ELISA titers, surface reactivity of vaccine-induced antibodies and the CSA binding inhibitory activity at CS2 pre-infection.**

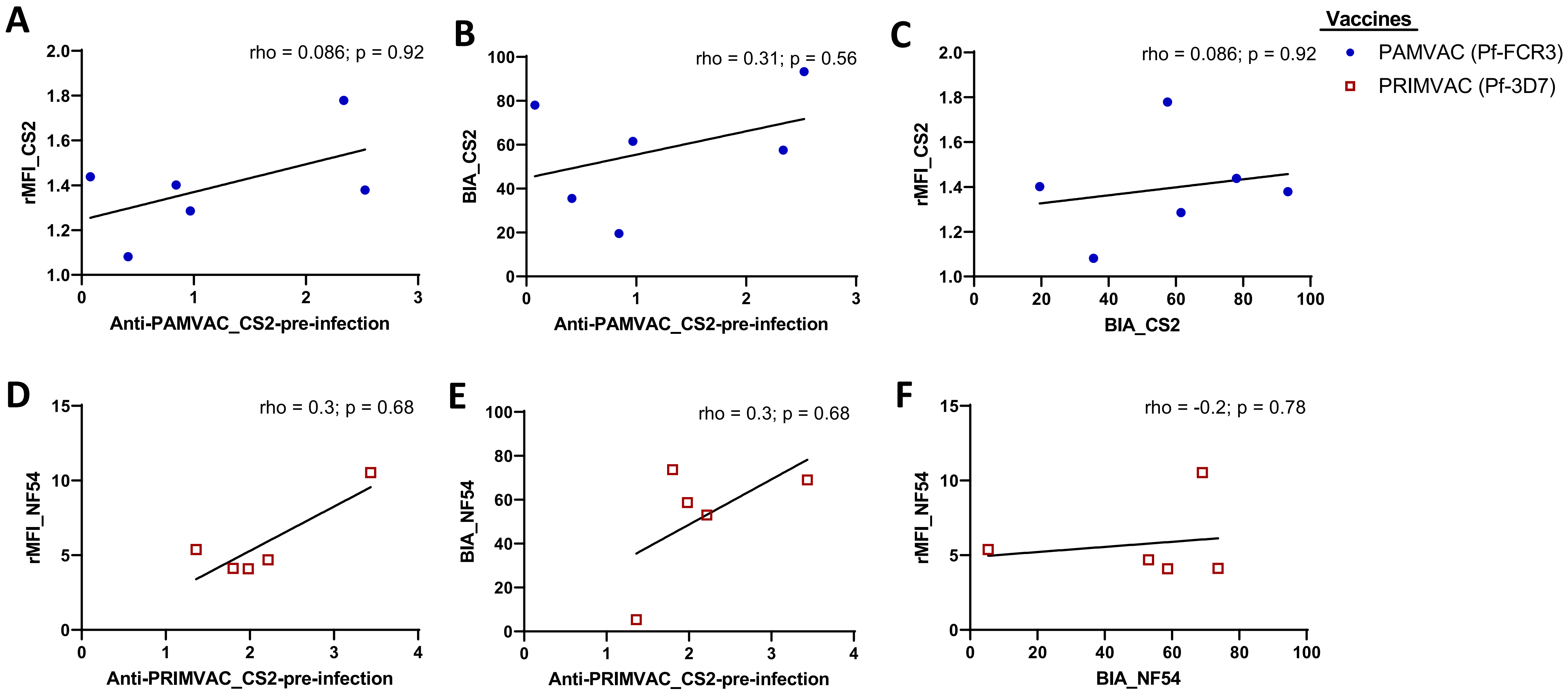

**Supplemental Figure S5: Correlations between ELISA titers, surface reactivity of vaccine-induced antibodies and the CSA binding inhibitory activity at 4 weeks postpartum.**

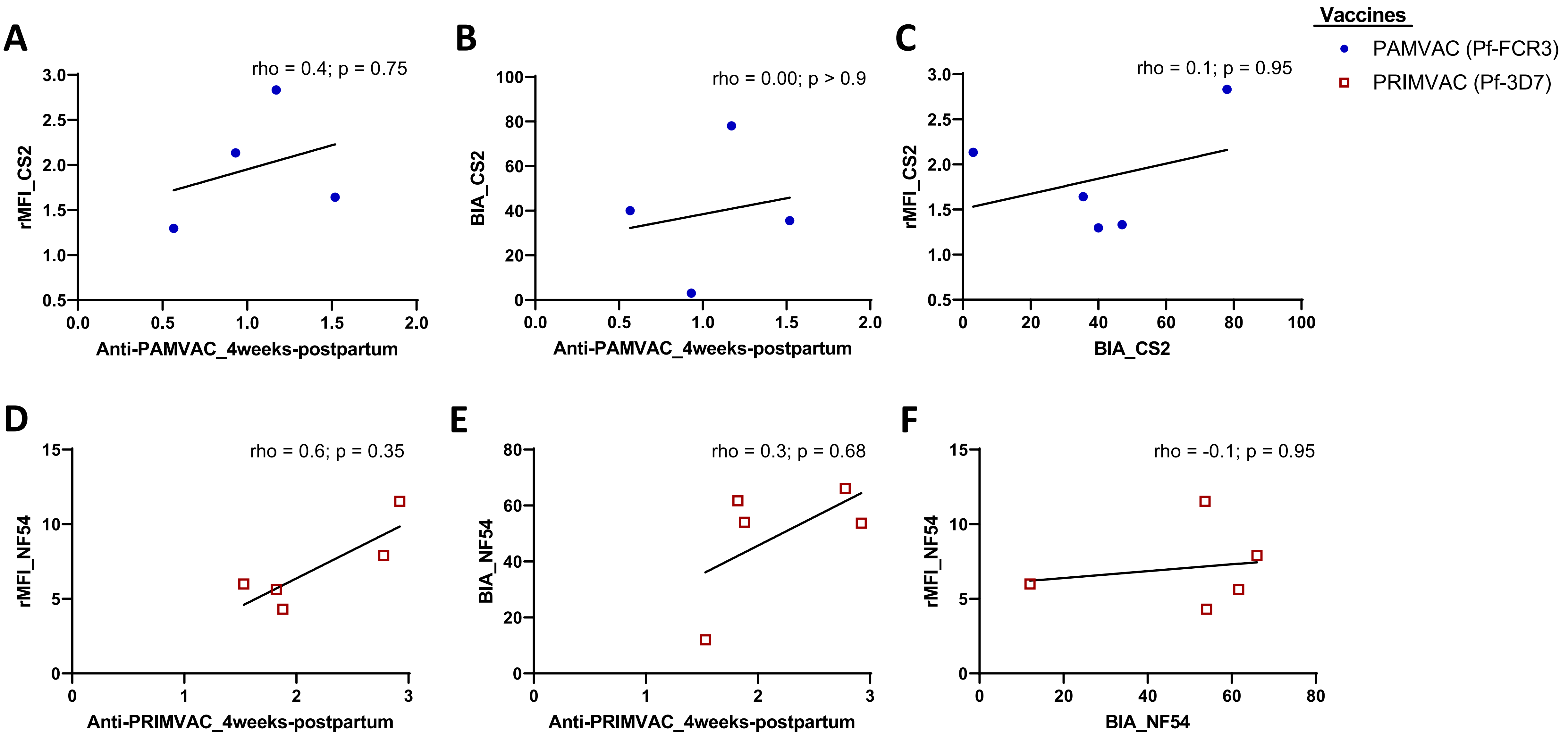

**Supplemental Figure S6: ELISA titer of PMV-induced antibodies in monkeys at D70, before CLP- or EPA-conjugated antigen boost, D14 and D56 post- boost.**

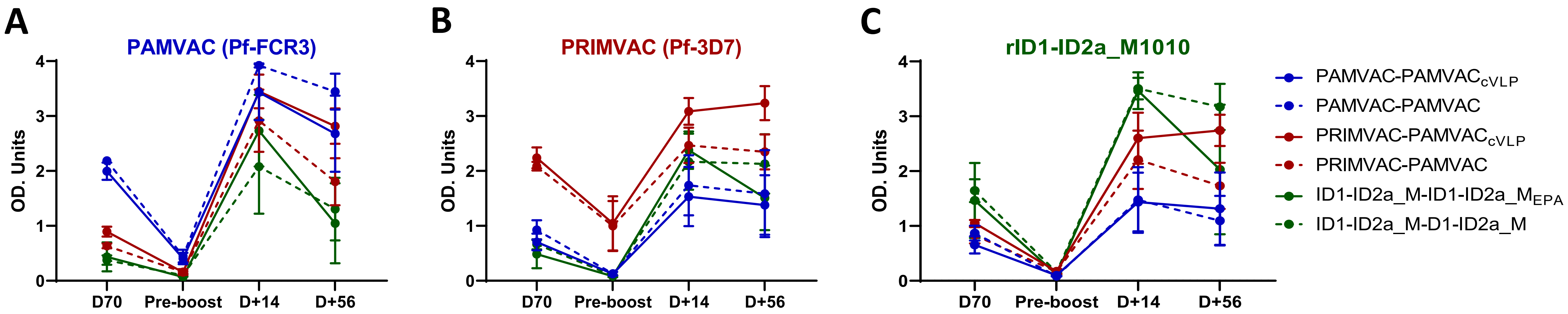

### Supplemental Figure S7: Peripheral and Placental Parasitemia detected by thin blood smears

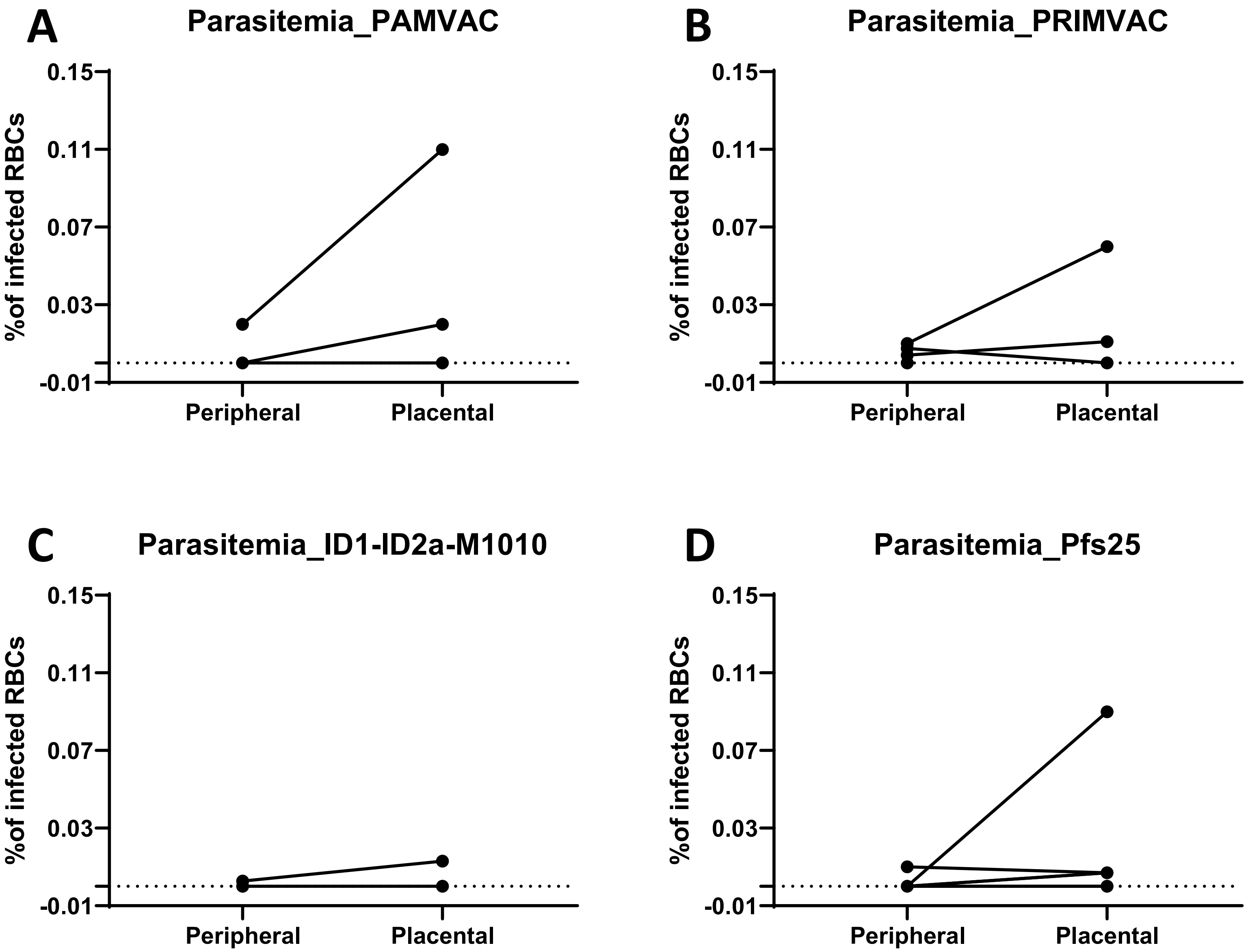

**Table S1: Binding characteristic for lots of CS2 inoculums**

| | No. of Vials | Binding phenotype (CD36/CSA) * |
|---|---|---|
| PfCS2 Lot-2 | 29 | 38/06 |
| PfCS2 Lot-3 | 26 | 49/02 |

**Note**: * Average bound infected red blood cells from 20 fields of 100X microscope, Binding receptors: CD36 and CSA
